# *(R)*-1-hydroxy-2-aminoethylphosphonate ammonia-lyase, a new PLP-dependent enzyme involved in bacterial phosphonate degradation

**DOI:** 10.1101/2020.12.13.422544

**Authors:** Erika Zangelmi, Toda Stanković, Marco Malatesta, Domenico Acquotti, Katharina Pallitsch, Alessio Peracchi

## Abstract

Phosphonates contain a particularly stable carbon-phosphorus bond, yet a number of microorganisms possess pathways to degrade these molecules and use them as source of phosphorus. One example is the widespread hydrolytic route for the breakdown of 2-aminoethylphosphonate (AEP). In this pathway, the aminotransferase PhnW initially converts AEP into phosphonoacetaldehyde (PAA), which is then cleaved by the hydrolase PhnX to yield acetaldehyde and phosphate.

This work focuses on a novel enzyme (hereby termed PbfA), which is often encoded in bacterial gene clusters containing the *phnW-phnX* combination. Although PbfA is annotated as a transaminase, we report that it catalyzes an elimination reaction on the naturally occurring compound *(R*)-1-hydroxy-2-aminoethylphosphonate (*R-*HAEP). The reaction releases ammonia and generates PAA, which can be subsequently hydrolyzed by PhnX. Overall, the PbfA reaction represents a frequent accessory branch in the hydrolytic pathway for AEP degradation, which expands the scope and versatility of the pathway itself.

---

Phosphonate compounds, containing a direct C–P bond instead of the more usual C–O–P ester linkage^1^, occur in the environment where they constitute both an important source of organic phosphorus^2^ and frequent environmental pollutants^3^. Phosphonates are difficult to degrade due to their very stable C-P bond, yet several microorganisms possess biochemical pathways that can break down some phosphonates, allowing their utilization as a source of phosphorus ^4,5^. Some of these pathways have been characterized at the biochemical and molecular level and can be schematically distinguished based on the mechanism through which the C-P bond is ultimately cleaved – either through a hydrolytic, radical or oxidative reaction^1,6^.

The degradation of phosphonates is particularly advantageous for microbes living in marine environments, where bioavailability of phosphorus is often a limiting factor for growth^7^. In particular, many marine bacteria possess a specialized ‘hydrolytic’ pathway for the breakdown of the natural compound 2-aminoethylphosphonate (AEP; also known as ciliatine), which is the most widely distributed biogenic phosphonate in the environment^4,8^. AEP degradation relies on an initial transamination catalyzed by PhnW, a pyridoxal 5’-phosphate (PLP) dependent aminotransferase, that converts AEP into phosphonoacetaldehyde (PAA)^9^; in turn, PAA is transformed into acetaldehyde and inorganic phosphate by the hydrolase PhnX (Figure 1A). In a variation of the above pathway, the PhnW-produced PAA can be converted first to phosphonoacetate and then to acetate and phosphate by the combined action of two other enzymes, termed PhnY and PhnA^10,11^ (Figure 1B). Degradation of another common aminophosphonate, L-phosphonoalanine (L-PA), proceeds quite similarly. It begins with the transamination of L-PA to phosphonopyruvate, operated by the PLP-dependent enzyme PalB; then a metal-dependent hydrolase, PalA, cleaves the phosphonopyruvate C–P bond to yield pyruvate and phosphate^4,12^ (Figure 1C).

**Figure 1.**
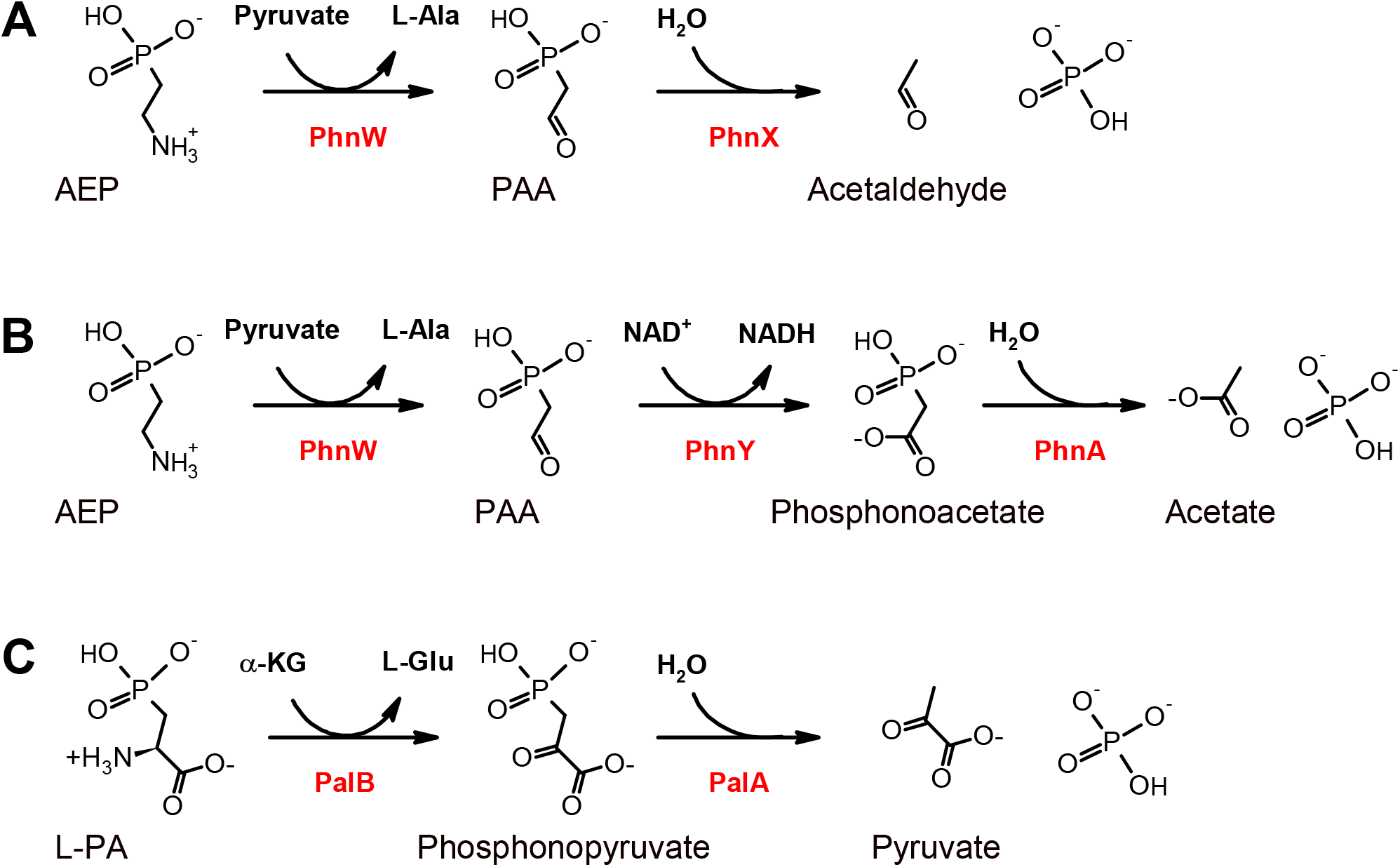
Known ‘hydrolytic’ pathways for the microbial catabolism of aminophosphonates, involving PLP-dependent enzymes. (A) AEP degradation: the transaminase PhnW converts AEP to PAA, which in turn is cleaved into acetaldehyde and phosphate by PAA hydrolase (PhnX). (B) A variant of the pathway above, in which PAA is converted to phosphonoacetate by PAA dehydrogenase (PhnY) and finally to acetate and phosphate by phosphonoacetate hydrolase (PhnA). (C) L-PA degradation: the amino acid is first transaminated by PalB, then the product phosphonopyruvate is cleaved by the hydrolase PalA into pyruvate and phosphate.

As shown in Figure 1, PLP-dependent enzymes play a pivotal role in different routes of aminophosphonate degradation. Since it is certain that additional enzyme systems (beyond those characterized so far) exist for the catabolism of phosphonates, we performed a detailed analysis of the genomes of many microorganisms that degrade phosphonates, looking in particular for genes that code for PLP-dependent enzymes and are associated to gene clusters for aminophosphonate breakdown.

This analysis led us to one gene (herein termed *pbfA*) that is often found within clusters containing the *phnW-phnX* (or *phnW-phnY-phnA*) combination. The *pbfA* gene product is usually annotated as a *γ*-aminobutyrate transaminase (GABA-T; Figure 2). While such an annotation appeared dubious, the gene’s involvement in phosphonate catabolism was strongly supported by the genomic context.

**Figure 2.**
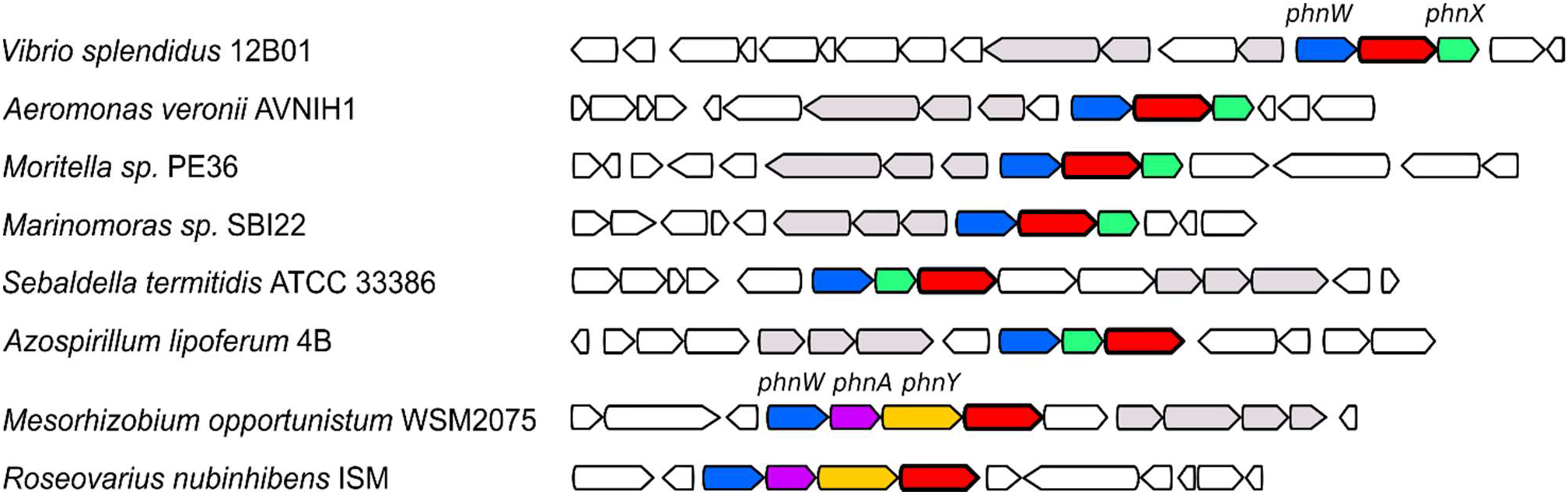
Recurring presence of a putative GABA-T gene (red) in clusters containing *phnW* (blue) and *phnX* (green) and in clusters containing *phnW, phnA* (purple) and *phnY* (yellow). Phosphonate-related transporter genes are shown in gray.

We therefore recombinantly produced the enzyme from the marine bacterium *Vibrio splendidus*, and assessed its activity against some potential substrates. Our results indicate that PbfA is a lyase acting on *(R)*-1-hydroxy-2-aminoethylphosphonate (*R-*HAEP, a compound related to AEP that occurs in nature^13–15^). The reaction catalyzed by PbfA generates PAA, which can be subsequently processed by PhnX.

## METHODS

The materials and experimental procedures used in this work are provided in the Supporting Information file.

## RESULTS AND DISCUSSION

### Bioinformatic identification of a putative transaminase involved in phosphonate degradation (PbfA)

We analyzed the genomic contexts of genes coding for proteins in the C-P hydrolase pathway, beginning with the AEP aminotransferase PhnW. Since it is known that PhnW can play a role both in the biosynthesis and in the degradation of AEP^10^, we focused on PhnW homologs (> 35% identical to the validated enzyme from *Salmonella*^9^) whose genes clustered either with *phnX* or with the duo *phnA/phnY* (*phnWX* and *phnWAY* clusters). Vice versa, we excluded cases in which *phnW* clustered with genes involved in phosphonate biosynthesis (*pepM* and *ppd*). By doing so we noticed that, in many bacteria, *phnW* and *phnX* were associated with a gene coding for a PLP-dependent enzyme, annotated as putative 4-aminobutyrate aminotransferase (GABA-T). This clustering occurred particularly in *γ* -proteobacteria belonging to the *Vibrionales, Aeromonadales, Alteromonadales* and *Oceanospirillales* orders. Furthermore, in some α-proteobacteria (such as in *Roseovarius nubinhibens*) very similar genes were found associated to the *phnWAY* operon (Figure 2). Due to the recurrent association of this GABA-T-like gene with clusters dedicated to AEP degradation, we provisionally labelled the encoded protein “phosphonate breakdown factor A” (PbfA).

For homology searches, we used the sequence of PbfA from *V. splendidus* 12B01 (EAP92040) as a query. When BLASTed against the PDB database of structurally characterized proteins, the *Vibrio* sequence showed the greatest similarity to enzymes such as GABA-T from *Escherichia coli* (1SF2; 30% identity) or β-alanine:pyruvate aminotransferase from *Pseudomonas aeruginosa* (4B98; 29% identity). These enzymes belong to a subfamily of PLP enzymes (called AT-II) that generally act on substrates whose amino group is not adjacent to a carboxylate^16^.

The vast majority of functionally characterized AT-II enzymes are aminotransferases, even though the subfamily includes at least one decarboxylase (2,2-dialkylglycine decarboxylase; DGD^17^), and a couple of enzymes that catalyze 1,2 eliminations (in particular, ethanolamine-phosphate phospho-lyase; ETNPPL^18^).

The similarity of PbfA to β-alanine aminotransferase could be compatible with PbfA catalyzing the transamination of AEP, which is a structural analog of β-alanine; however, this would be an unneeded duplicate of the reaction catalyzed by PhnW. This argument suggested that PbfA must act on a compound different from AEP and/or catalyze a reaction different from transamination. An alignment of the PbfA sequences with those of other AT-II enzymes showed mutations at some key residues known to be important for substrate and reaction specificity (Supplemental Results and Fig. S1).

Hypotheses on the possible activity of the enzyme were developed based on the following considerations: (a) the substrate of PbfA must necessarily contain a primary amino group, as is the norm with PLP-dependent enzymes; (b) the substrate is presumably a phosphonate compound rather abundant in nature, to justify the recurrence of the PbfA gene; and (c) the product of the reaction should be either AEP or PAA, in order to feed into the PhnW-PhnX pathway. Based on these considerations, two possibilities seemed the most convincing. The first was that PbfA could be producing AEP from L-PA, through a decarboxylation reaction analogous to that catalyzed by DGD. A second possibility was that PbfA could catalyze a 1,2-elimination on 1-hydroxy-2-aminoethyl phosphonate (HAEP, a compound similar to AEP found in different organisms^13,19^), to directly generate PAA – a reaction somewhat similar to that of ETNPPL.

Examination of the complete genomes of the bacteria possessing PbfA showed that they almost invariably lacked the known enzymes for L-PA breakdown (PalB and PalA, Figure 1) and none of them possessed the only known enzyme for HAEP degradation (the oxidase PhnZ^15^).

### PbfA is neither a transaminase nor a decarboxylase, but a lyase acting on *R*-HAEP

To test the function of PbfA, we recombinantly produced the enzyme from *V. splendidus*, together with PhnW and PhnX from the same organism. We then assessed the activity of purified PbfA *in vitro* against the potential substrates.

First we tested a possible reaction of PbfA with AEP. When the enzyme was incubated with AEP for several hours, no consumption of the aminophosphonate was observed by TLC. Similarly, when the same experiment was conducted in the presence of an amino group acceptor such as glyoxylate, pyruvate or α-ketoglutarate, no formation of new amino acids was observed, implying that PbfA, as predicted, is unable to transaminate AEP. This was confirmed by the fact that no formation of acetaldehyde was detected in a reaction mixture where PbfA was incubated with AEP plus amino group acceptors, PhnX, NADH and alcohol dehydrogenase (ADH).

Additionally, when PbfA was incubated with D,L-PA under the same conditions as above, no reaction was detected either by TLC or by a coupled assay with PhnW, PhnX and ADH. These results ruled out the possibility that PbfA could act as a decarboxylase (or as a decarboxylation-dependent transaminase, like DGD) on phosphonoalanine, to generate AEP or PAA.

In contrast to the results summarized above, we observed that incubation of PbfA with racemic HAEP led to the release of ammonia, as expected in a 1,2-elimination reaction (Figure 3A). The generation of PAA (the other postulated product of the reaction) was inferred by coupling the PbfA reaction with PhnX and ADH (Figure 3A). Phosphate was not released in the reaction of PbfA with HAEP, except when PhnX was also present (BIOMOL Green assay, data not shown). The PbfA-catalyzed reaction consumed only about 50% of the racemic substrate, implying that only one of the two HAEP enantiomers was used by the enzyme (Figure 3B).

**Figure 3.**
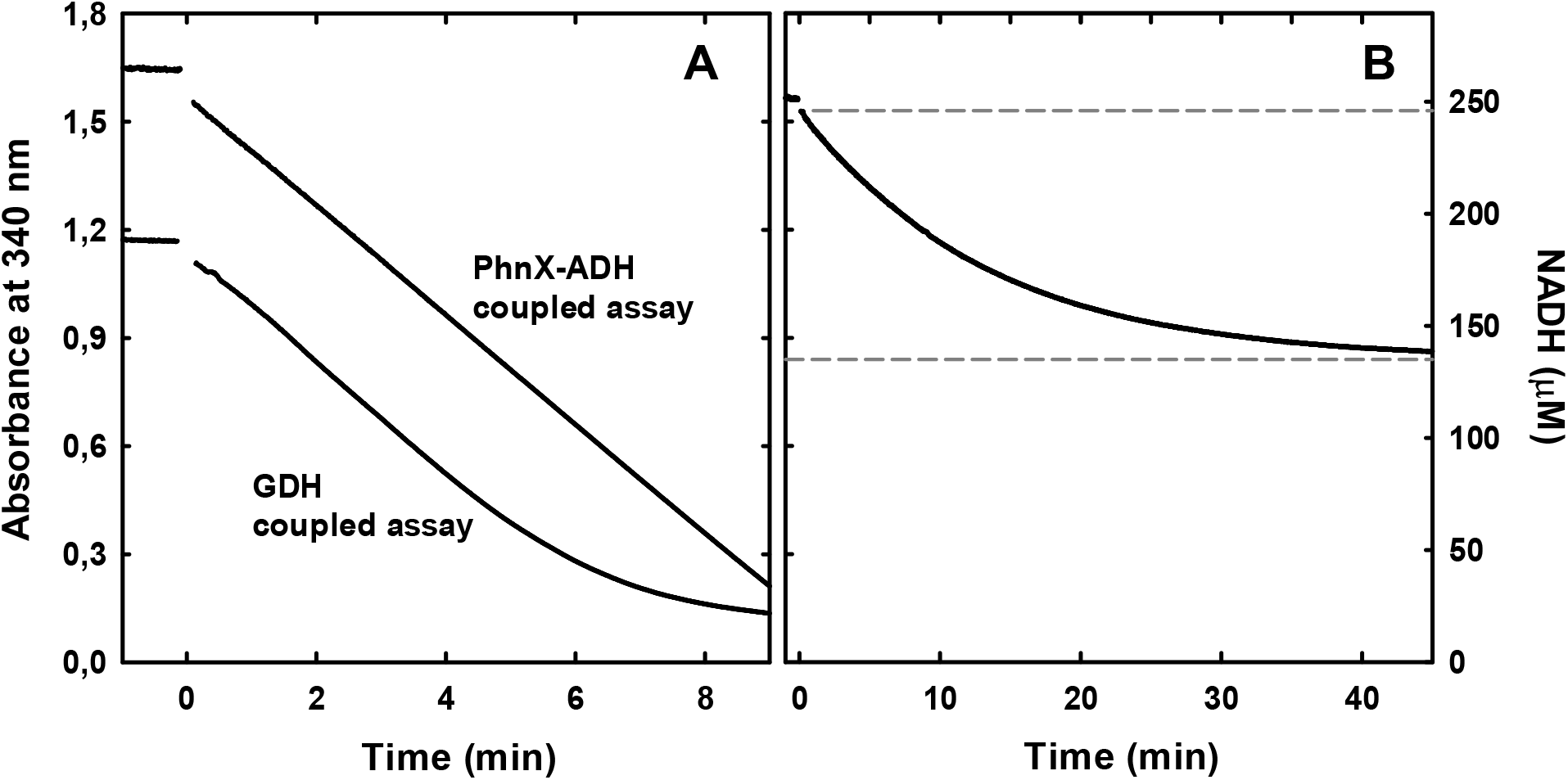
(A) Coupled assays to monitor the reaction of PbfA with HAEP. The assay with GDH (lower curve) monitored the release of ammonia. The reaction contained 10 mM racemic HAEP, 1 mM α-ketoglutarate, ∼0.25 mM NADH, 5 mM MgCl_2_, 5 µM PLP, in addition to PbfA (2.9 µM) and GDH, in triethanolamine-HCl pH 8.0. The coupled assay with PhnX and ADH (upper curve) detected the ultimate formation of acetaldehyde. Reaction conditions were as above, except that α-ketoglutarate and GDH were omitted while PhnX and ADH were included. The very similar slopes obtained in the two assays are consistent with the two processes being both rate-limited by the same step, namely the elimination of HAEP. (B) Coupled assay with PhnX and ADH conducted as above, but in the presence of only 0.2 mM racemic HAEP (and 4.9 µM PbfA). Based on NADH consumption, just about 0.1 mM HAEP was used by PbfA, strongly suggesting that only one of the two HAEP enantiomers is a substrate.

To understand the reaction specificity of the lyase, we synthesized the two HAEP enantiomers as described^20^ and tested them individually for reaction with PbfA. These experiments showed that *R-*HAEP is efficiently processed by the enzyme, whereas *S-* HAEP is completely unreactive. When the reaction between PbfA and *R-*HAEP was monitored by ^1^H-NMR, a consumption of this aminophosphonate was observed, with the simultaneous appearance of a compound showing the NMR signature of PAA^21^ (Figure 4).

**Figure 4.**
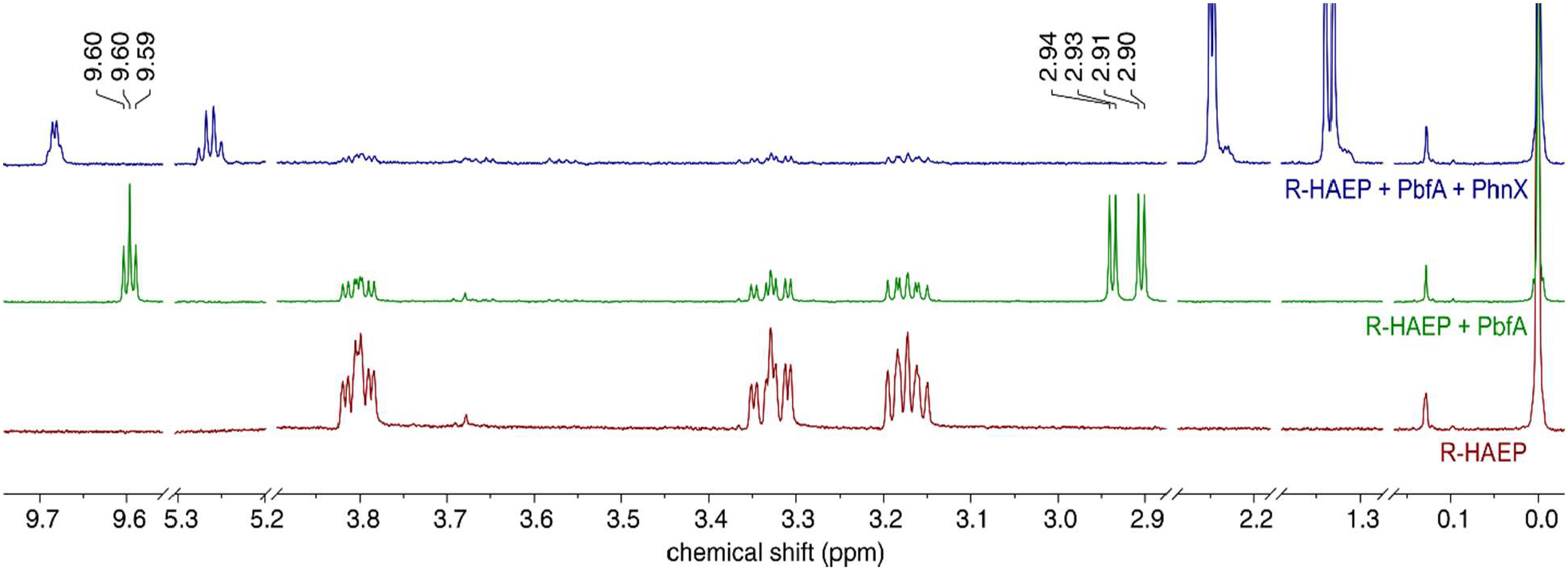
1H-NMR spectra of *R*-HAEP before (bottom, red line) and after (middle, green line) a 15-min incubation with PbfA. The new peaks at 2.90-2.94 ppm and at 9.6 ppm are attributed to PAA based on published data^21^ and on the direct comparison to the spectrum of PAA generated upon transamination of AEP by PhnW (see Supplementary Figure S4). Addition of PhnX (top, blue line) led to disappearance of the peaks mentioned above and to the appearance of new peaks attributable to acetaldehyde.

This proved that PbfA is a lyase acting on *R-*HAEP. The observed 1,2-elimination reaction is similar to those catalyzed by other PLP-dependent enzymes such as L-threonine ammonia-lyase (EC 4.3.1.19). In analogy with the catalytic mechanism previously suggested for various PLP-dependent lyases^22,23^, a tentative mechanism for the elimination reaction catalyzed by PbfA can be outlined (Figure S2).

### PbfA is a rather efficient and highly specific enzyme

As shown in Figure 3, we could monitor the kinetics of *R-*HAEP elimination by two different coupled assays. However, the GDH-based assay, despite requiring only one coupling enzyme, was characterized by an initial lag phase very hard to eliminate^24^, presumably due to the complex allosteric behavior of GDH^25^. Consequently, for the kinetic characterization of PbfA we used the coupled assay with PhnX and ADH.

At pH 8.0 and 25°C, the observed kinetic parameters were: k_cat_ = 5.3 ± 0.03 s^-1^, K_M_ = 0.36 ± 0.04 mM, and k_cat_/ K_M_ =14600 M^-1^s^-1^ (Figure S3). These values are comparable to, or better than, those reported for other PLP-dependent lyases such as ETNPPL^18^. They might even be underestimates, since we found that the specific activity of our purified PbfA stocks, once thawed, tended to decrease significantly within a few hours.

Tests on some commercially available HAEP analogs showed that the lyase activity of PbfA is very specific for HAEP. When the PO_3_ group of HAEP was absent (as in ethanolamine) or replaced by a carboxylate (as in D,L-isoserine, i.e., 3-amino-2-hydroxypropionate), and even if the OH group was substituted by other good leaving groups, such as bromine or a thiol (in bromoethylamine and cysteamine, respectively), the elimination reaction was virtually undetectable (data not shown).

These observations, together with the constant localization of *pbfA* in gene clusters for phosphonate breakdown, confirm that the physiological role of this enzyme is strictly dedicated to *R*-HAEP degradation. We hence propose for PbfA the systematic name *(R)*-1-hydroxy-2-aminoethylphosphonate ammonia-lyase.

### *R*-HAEP is not efficiently degraded by PhnW and PhnX

When we incubated the recombinant *V. splendidus* PhnW with its standard substrates, AEP and pyruvate, formation of the expected transamination products, alanine and PAA, was easily detected by ^1^H-NMR (Figure S4A). The reaction kinetics could be monitored trough the coupled assay with PhnX and ADH, yielding catalytic parameters (k_cat_= 15.5 ± 0.3 s^-1^, K_M_^AEP^=3.2 ± 0.15 mM, k_cat_/ K_M_^AEP^ = 4900 M^-1^s^-1^) comparable to those reported for the *Salmonella* PhnW^9^.

When *R-*HAEP was incubated with PhnW and pyruvate, in the absence of PbfA and PhnX, we could observe the formation of alanine by both TLC (ninhydrin staining) and NMR (Figure S4B), meaning that HAEP was getting transaminated to some extent. The ^1^H-NMR data showed decrease of the *R*-HAEP and pyruvate peaks and the appearance of new peaks for alanine, but also for an aldehydic proton, belonging to a compound different from both PAA and acetaldehyde (Figure S4B) which did not disappear upon the addition of PhnX.

When we assessed the consecutive reactions of PhnW and PhnX on *R*-HAEP, through a coupled assay with ADH, we did not observe any immediate NADH oxidation, implying that no aldehyde amenable to reduction by ADH was being produced. However, upon incubating *R*-HAEP with PhnW and pyruvate for 18 h, and finally adding NADH and a large amount of ADH, we could detect a modest (and rather slow) NADH oxidation, suggesting that some suitable ADH substrate had formed over time. Some small amount of phosphate was also released upon incubation of *R*-HAEP with PhnW, and this release was independent of the presence of PhnX (Figure S5). The total amount of phosphate released was much lower than the amount of alanine formed, implying that phosphate release, albeit dependent on the PhnW transamination reaction, is not tightly coupled to it.

Although we could not positively identify the aldehyde generated by the activity of PhnW on *R*-HAEP, our tentative interpretation of the data above is the following. We assume that the transamination of *R*-HAEP yields the expected transamination product *(R)*-1-hydroxy-phosphonoacetaldehyde, which however is not a substrate for either PhnX or ADH. Upon time, 1-hydroxy-phosphonoacetaldehyde breaks down to yield phosphate and glycolaldehyde (which is a substrate of ADH, albeit suboptimal). This ‘non-enzymatic’ breakdown of the C-P bond would be analogous to the reported spontaneous decarboxylation of tartronate semialdehyde, which also yields glycolaldehyde ^26,27^.

Irrespective of the actual mechanism, our results shown that *R-*HAEP, despite being structurally similar to AEP, cannot be properly processed through the PhnW-PhnX pathway, an observation that might justify the very existence of a dedicated degradative enzyme. Note that also the gene for the oxidase PhnZ, that converts *R*-HAEP to glycine and phosphate, is sometimes found associated with the *phnWX* or *phnWAY* clusters^6^.

## CONCLUSIONS

Through a combination of bioinformatics, enzymology and organic chemistry^28^, we have identified and functionally characterized a PLP-dependent enzyme, termed PbfA, which is generally annotated in public databases as GABA-T. Recurrence of *pbfA* in gene clusters dedicated to AEP degradation had been incidentally noted before^4,6^, but without any experimental follow-up. We have shown that PbfA from *Vibrio* (and hence presumably also its close structural and positional homologs found in many bacteria) catalyzes a very specific, and previously undescribed, elimination reaction on *R-*HAEP.

The PbfA reaction represents a recurrent accessory branch in the well-established hydrolytic AEP degradation pathway and serves to funnel *R-*HAEP into the PhnW-PhnX pipeline (Figure S6). The ability to consume *R-*HAEP, in addition to AEP, increases the flexibility of the hydrolytic pathway, presumably allowing a bacterium to exploit different phosphorus sources and hence increasing its fitness under variable nutrient availabilities.

*R-*HAEP was first isolated as a hydrolysis product of complex polysaccharides of the soil amoeba *Acanthamoeba castellanii* ^13,14^; furthermore, *R*-HAEP is an intermediate in yet another pathway for the degradation of AEP, based on the reactions of the two oxygenases PhnY* and PhnZ^15,29^. As the combination *phnW-phnX-pbfA* (or the nearly equivalent combination *phnW-phnY-phnA-pbfA*) is widespread and abundant among marine bacteria (Figure 2), *R-*HAEP may represent an important additional source of phosphorus in certain environments.

The function of the enriched hydrolytic pathway encompassing PhnW, PhnX and PbfA is analogous to that of the oxidative pathway composed the enzymes PhnY* and PhnZ^15,29^, in the sense that either pathway can obtain phosphate from both AEP and *R-* HAEP. One significant difference is that the PhnY*-PhnZ reactions, requiring molecular oxygen, are not going to be functional under anaerobic conditions, whereas the *phnW-phnX-pbfA* cluster is found also in some strict anaerobes.

## Supporting information

Supplementary Methods, results and figures

## Abbreviations

AEP: 2-aminoethylphosphonate;
HAEP: 1-hydroxy-2-aminoethylphosphonate;
L-PA: L-phosphonoalanine;
PAA: phosphonoacetaldehyde;
PhnW: 2-aminoethylphosphonate:pyruvate aminotransferase;
PhnX: phosphonoacetaldehyde hydrolase (phosphonatase);
PhnY: phosphonoacetaldehyde dehydrogenase;
PhnA: phosphonoacetate hydrolase;
PalA: phosphonopyruvate hydrolase;
PalB: phosphonoalanine aminotransferase;
GABA-T: 4-aminobutyrate aminotransferase;
RidA: 2-iminopropanoate deaminase;
ADH: alcohol dehydrogenase;
GDH: glutamate dehydrogenase;
AT-II: subgroup-II aminotransferases (also known as aminotransferases class III);
DGD: 2,2-dialkylglycine decarboxylase;
ETNPPL: ethanolamine-phosphate phospho-lyase;
PLP: pyridoxal-5′-phosphate.

## AUTHOR CONTRIBUTIONS

E.Z. performed the bioinformatic analysis, proteins purification, NMR measurements and biochemical studies. T.S. and K.P. synthesized the HAEP and contributed to design experiments. M.M. contributed to the bioinformatic analysis and NMR measurements. D.A. contributed to the NMR measurements. A.P. conceived and supervised the project and wrote the manuscript. All the authors have read and approve the final version of the manuscript. The authors declare no competing financial interest.

## ACKNOWLEDGEMENTS

This work has been supported in part by a grant from the “Fondazione Cariparma” (to A.P.). Research has also benefited from the framework of the COMP-HUB Initiative, funded by the ‘Departments of Excellence’ program of the Italian Ministry for Education, University and Research (MIUR, 2018-2022). K.P greatly acknowledges the financial support by the Austrian Science Fund (FWF): P27987-N28.

## SUPPORTING INFORMATION

The Supporting Information file includes the Supplementary Materials (experimental details about bioinformatics analyses, proteins purification, NMR measurements, spectrophotometric enzyme assay) Supplemental Results (analysis of the frequence of PbfA occurrence in bacteria; sequence comparison between PbfA and other, phylogenetically related, PLP-dependent enzymes) and figures S1-S6.

