## Supplementary Methods, results and figures for "*(R)*-1-hydroxy-2-aminoethylphosphonate ammonia-lyase, a new PLP-dependent enzyme involved in bacterial phosphonate degradation"

SUPPLEMENTAL MATERIALS AND METHODS

### Materials

Alcohol dehydrogenase (ADH, baker's yeast), glutamate dehydrogenase (GDH, beef liver) and DL-PA were from Sigma. 2-aminoethylphosphonate (AEP) was from Wako chemicals. Triethanolamine (TEA), pyridoxal-5'-phosphate (PLP), bovine serum albumin (BSA) were from Sigma-Aldrich. NADH was from Alfa Aesar. TLC plates (silica gel 60 $F_{254}$) were from Merck Millipore. Deuterium oxide was from VWR International and deuterated trimethylsilyl propionic acid (TSP) from Stohler Isotope Chemicals. All other reagents were from Fluka or Sigma–Aldrich.

Racemic HAEP was synthesized beginning from vinylphosphonic acid, as described in [1]; the *(S)*- and *(R)* enantiomers of HAEP were also prepared by methods described in the same publication[1].

### Bioinformatics analyses

Searches for gene homologs in complete genomes were performed by using BlastP [2] on the Assembly database of NCBI (https://www.ncbi.nlm.nih.gov/assembly/) and on the website of the Department of Energy, Integrated Microbial Genomes & Microbiomes (IMG/MER) (https://img.jgi.doe.gov/cgi-bin/mer/main.cgi). Visualization of genomic contexts for bacterial genes was also performed using tools available on IMG/MER and on the MicrobesOnline website (http://www.microbesonline.org/). Protein similarity to functionally validated PLP-dependent proteins was assessed by consulting the B6 database [3]. Multiple sequence alignments were created with ClustalX2 and displayed with ESPript 3.0[4].

### Plasmid construct and protein purification

The *Vibrio splendidus* 12B01 genes for PhnW (GenBank: EAP92041), PhnX (GenBank: EAP92039) and PbfA (GenBank: EAP92040) were codon-optimized for expression in *Escherichia coli*, synthesized and inserted into pET24a-C-HIS, pET28a-N-HIS and pBluesScript II KS(+), respectively, by BaseClear BV (Leiden, the Netherlands).

The *pbfA* gene was later subcloned into a pET28-CpoI expression vector[5] exploiting the CpoI restriction site, so as to obtain the expression of a N-terminal His6-tagged protein. The *E. coli* XL1-Blue strain was used for cloning applications and plasmid amplification. The subcloned *pbfA* gene was sequence-verified and the plasmid was used to transform *E. coli* Tuner cells (EMD Biosciences) for protein expression. PhnX was also overexpressed in Tuner cells whereas PhnW was expressed in *E. coli* BL21 star cells.

Cultures were grown at 37 °C in LB medium supplemented with kanamycin (50 μg/mL) until $OD_{600}$ reached ~0.7, at which point the temperature was reduced to 20 °C and protein production was induced by adding 0.3 mM isopropyl-β-D-1-thiogalactopyranoside (IPTG). 20 hours after induction, the cells were harvested by centrifugation, pellets were resuspended in an appropriate lysis buffer, as follows: PhnW and PbfA: 50 mM phosphate buffer pH 7.5, 150 mM NaCl, 5 mM β-mercaptoethanol, 1 mM PMSF, 1 mM benzamidine, 1 mg/ml lysozyme, 10 μM PLP; PhnX: 50 mM HEPES pH 7.5, 200 mM NaCl, 5 mM β-mercaptoethanol, 1 mM PMSF, 1 mM benzamidine, 1 mg/ml lysozyme, 10 mM $MgCl_2$.

Then the cells suspensions were sonicated and centrifuged. Cleared cell lysates were loaded on a His-Select® cobalt affinity resin (Sigma-Aldrich) and the recombinant proteins were purified following the manufacturer's instructions. Protein purity was assessed by SDS-PAGE. Fractions with purity higher than 90% were pooled and dialyzed against storage buffer.

PhnW: 50 mM Na-phosphate pH 7.5, 300 mM NaCl, 1 mM DTT, 5 μM PLP;

PhnX: 50 mM HEPES pH 7.5, 1 mM DTT, 10 mM $MgCl_2$;

PbfA: 50 mM HEPES pH 7.5, 150 mM NaCl, 1 mM DTT, 5 μM PLP).

After dialysis, 10% (v/v) glycerol was added to the enzyme stocks and their concentration was assessed spectrophotometrically. Each stock was subdivided in aliquots (about 0.5 ml each), which were frozen and stored at -80°C.

Protein concentrations were determined using the following $\varepsilon_{280nm}$ values, estimated from the amino acid sequence by the ProtParam tool (http://web.expasy.org/protparam/) 48360 $M^{-1}$ $cm^{-1}$ for PhnW, 32430 $M^{-1}$ $cm^{-1}$ for PhnX and 46870 $M^{-1}$ $cm^{-1}$ for PbfA. The protein yield was 22 mg, 50 mg and 47 mg per liter of culture for PhnW, PhnX and PbfA respectively.

**Qualitative detection of enzyme reactions by thin layer chromatography (TLC)**

For the qualitative detection of enzyme activity, PhnW and PbfA were incubated at 37 °C in 50 mM TEA-HCl buffer (pH 8.0), in the presence of substrates at the desired concentrations. At the end of the incubation, two microliters of the reaction mixture were spotted on a thin-layer silica gel plate and developed (acetic acid:1-propanol:distilled water - 1:3:1). At the end of the chromatographic separation, amino group-containing compounds were visualized by spraying the plates with 0.3% ninhydrin in methanol, drying and heating for about 5 min.

**Measurement of phosphate release**

Phosphate release during the course of enzymatic reactions was assessed by using the BIOMOL® Green kit (Enzo Life Sciences), according to the manufacturer's instructions. The reactions mixture contains 50 mM TEA-HCl buffer pH 8.0, 1 mM DTT, 5 µM PLP, 100 mM KCl, 5 mM $MgCl_2$, 2 µM PhnW or PbfA, 2 µM PhnX, 1 mM of AEP or R-HAEP and 1 mM pyruvate (if specified). The reactions were carried out at room temperature and they were stopped after 1 hour by adding a 12-fold excess of BIOMOL® Green. Color development was read 30 min later measuring $OD_{620nm}$ on a Cary 50 UV-Vis spectrophotometer (Varian).

**NMR measurements**

$^1$H NMR spectra were collected with a JEOL ECZ600R spectrometer in non-spinning mode at 25°C using Simple DANTE presat sequence for $H_2O$ suppression. The reaction mixture contained 50 mM potassium phosphate pH 8.0, 100 mM KCl, 5 mM $MgCl_2$, 5 mM of substrates (AEP or *R*-HAEP, and pyruvate if needed), enzymes (0.8 µM PhnW or 0.8 µM PbfA, 1 µM PhnX) and 1 mM of deuterated TSP, used as internal chemical shift reference (δ=0.00 ppm), in 450 µl $H_2O$ plus 50 µl $D_2O$. NMR spectra were processed and analyzed with MestReNova v12.0.4 (Mestrelab Research).

**Spectrophotometric enzyme assays**

The PhnW-catalyzed transamination of AEP (with pyruvate as a co-substrate) was measured through a coupled assay with PhnX and ADH [6]. Solutions for the coupled assay (150 µl final volume) typically contained 50 mM buffer (TEA-HCl) pH 8.0, 0.5 mg/ml BSA, ~0.25 mM NADH, 5 mM $MgCl_2$, 5 µM PLP, in addition to the enzymes and substrates. Unless otherwise indicated, reactions were carried out at 25°C and started by the addition of 0.13 µM PhnW. The reaction was measured by monitoring the disappearance of NADH at 340 nm on a Cary 400 thermostatted spectrophotometer (Varian).

The PbfA-catalyzed elimination of *R*-HAEP was also monitored by coupling this reaction with the reactions of PhnX and ADH, or *via* a coupled assay with glutamate dehydrogenase (GDH). In both cases, the reaction was followed by monitoring the disappearance of NADH at 340 nm.

The coupled assay with PhnX and ADH was typically conducted in TEA-HCl buffer, pH 8.0. The reaction mixture also contained ~0.25 mM NADH, 5 mM $MgCl_2$, 5 μM PLP, PhnX and ADH, in addition to variable amounts of PbfA and of *R*-HAEP.

The coupled assay with GDH, detecting the release of ammonia, was conducted under the same conditions as described above, except that PhnX and ADH were omitted, while the reaction mixture contained 1 mM α-ketoglutarate and 6.7 U GDH.

Kinetic data were analyzed by nonlinear least-squares fitting to the appropriate kinetic equation (e.g., the Michaelis-Menten equation) using Sigma Plot (Systat Software Inc.).

SUPPLEMENTAL RESULTS

**Sequence analysis of PbfA from *Vibrio splendidus* 12B01**

As stated in the main text, PbfA belongs to a structural subgroup of PLP-dependent enzymes, termed “AT-II” (or “class III aminotransferases”)[7]. Most enzymes in this subgroup are aminotransferases. Furthermore, nearly all enzymes in this subgroup act on substrates whose amino group is not adjacent to a carboxylate[7].

Through a multiple sequence alignment with functionally and structurally characterized AT-II enzymes, we could identify in the sequence of PbfA from *Vibrio splendidus* 12B01 the catalytic lysine (K 317) and other conserved residues that interact with the PLP cofactor (T347, G144 and T145, which bind the phosphate group; D288, which interacts with the pyridine nitrogen and F171, which stacks onto the PLP ring) (Figure S1). Furthermore, we examined the active site residues that, in AT-II aminotransferases, differentiate those enzymes using α-ketoglutarate from those that use pyruvate as an amino group acceptor. In the case of α-ketoglutarate-specific transaminases, the active site contains an arginine (R398 in GABA-T from *E. coli*) which forms a salt bridge with the α-carboxylate of the ketoacid. When the acceptor is pyruvate, the corresponding arginine is at a different position (Figure S1). Also, in proteins that use α-ketoglutarate there is a conserved glutamate (211 in GABA-T from *E. coli*) which is involved the 'Gateway system' (E211 binds R398 in the first half-reactions, preventing transamination of α-amino groups instead ω-amino groups, and releasing it in the second part allowing the correct positioning of the keto acid’s α-carboxylate) and another arginine (R 141 GABA-T from *E. coli*) which interacts with the γ-carboxylate of α-KG[7].

In case the substrate does not have the carboxyl group in place of arginine there are other specific residues for interaction. For example in the lyase ETNPPL it is replaced by a histidine and in 2,2-dialkylglycine decarboxylase (DGD) by a methionine. PbfA lacks the two arginines and glutamate typical of AT-II transaminases using the α-KG as the amino group acceptor. In particular, arginine 141 of GABT_ECOLI is consistently replaced by an alanine (A174 in the enzyme from *V. splendidus*). In a sense, PbfA is more similar to β-alanine-pyruvate transaminase, which contains a hydrophobic amino acid (V 155 in the enzyme from *Pseudomonas putida*) at this position. There is also a conserved lysine (K 438 of PbfA from *V. splendidus*) in the same position where we find the arginine in β-alanine-pyruvate transaminase and in other AT-II transaminases that use pyruvate as an amino group acceptor (Figure S1).

### SUPPLEMENTAL FIGURES

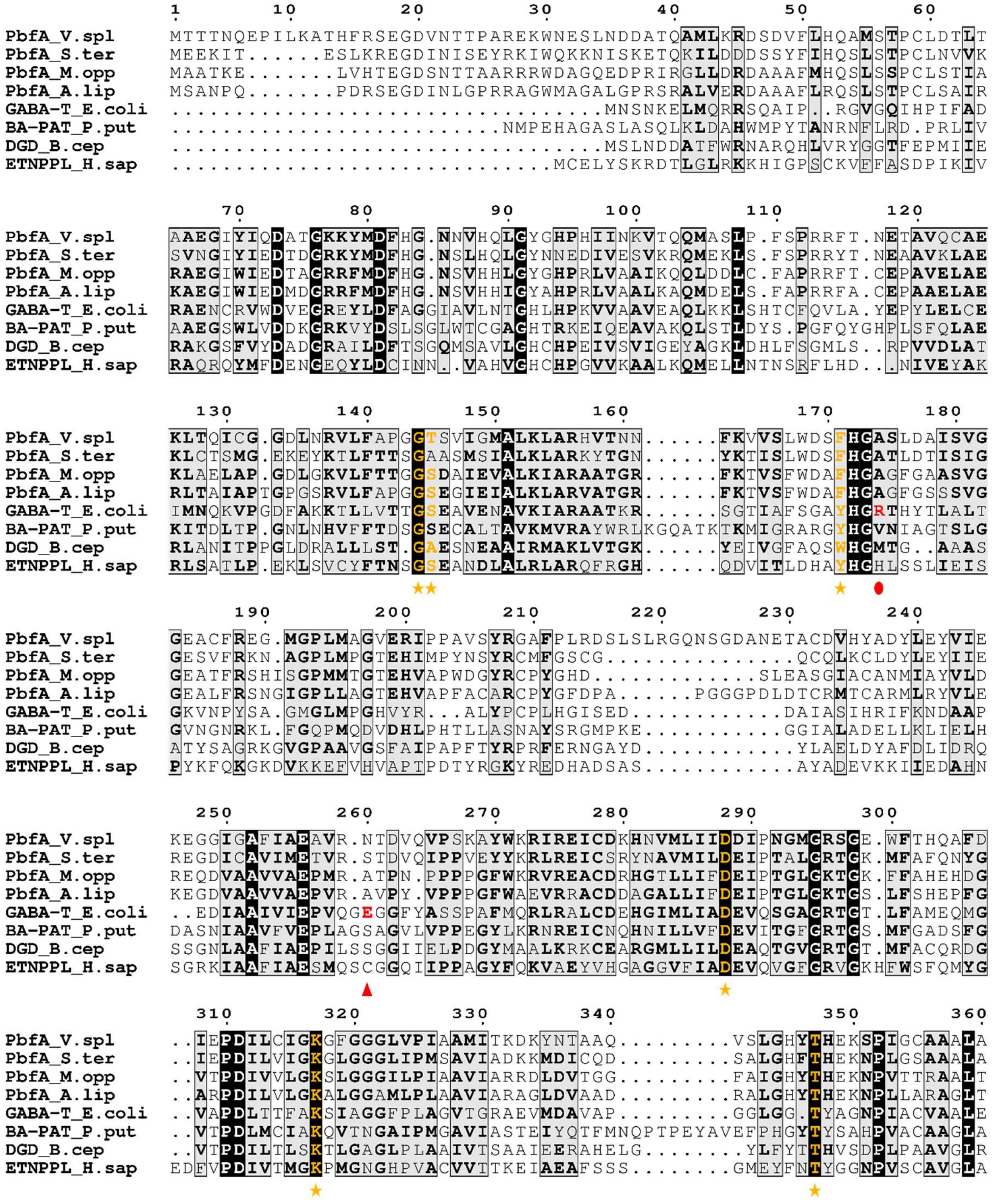

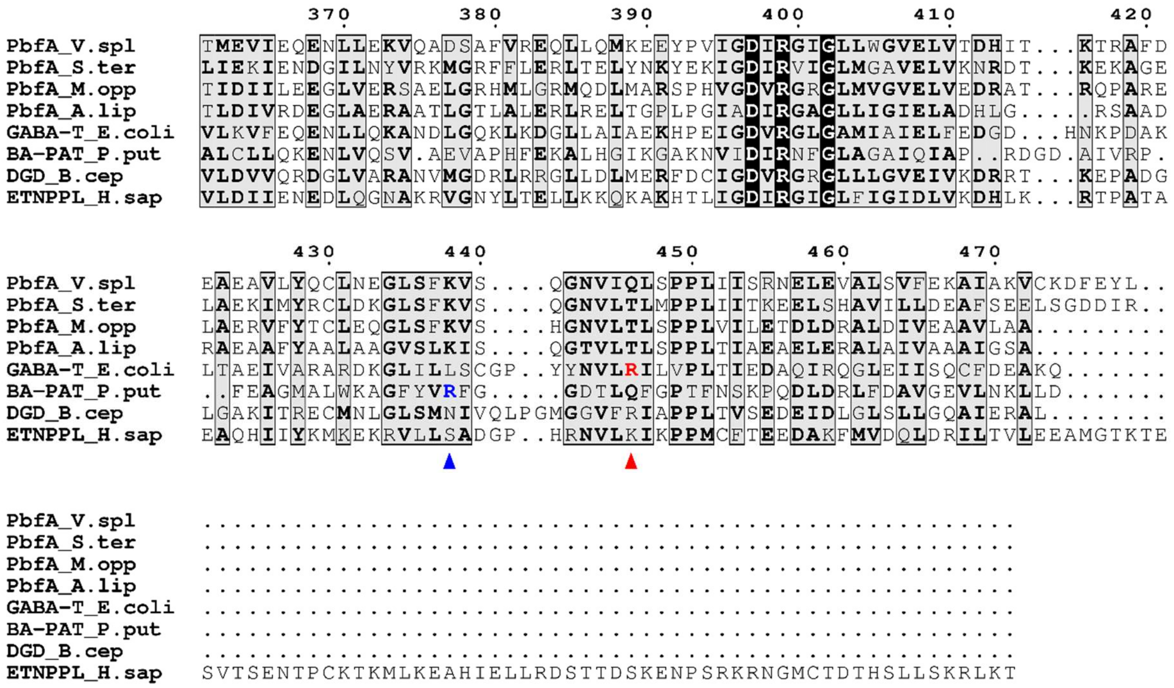

**Figure S1** – Multiple sequence alignment of four PbfA proteins (corresponding to genes signaled as hypothetical GABA-T in Figure 2 of the main text) with functionally validated enzymes belonging to the same subgroup of PLP-dependent enzymes (AT-II), namely GABA-T from *E. coli* (NP_417148.1), DGD from *Burkholderia cepacia* (P16932.3), ETNPPL from *Homo sapiens* (NP_112569.2 ) and β-alanine:pyruvate transaminase from *Pseudomonas putida* (BA-PAT, P28269.1). The orange stars indicate residues interacting with PLP and the catalytic lysine residues (K 317 in PbfA from *Vibrio splendidus*, PbfA_V.spl). Red symbols point to the residues typical of aminotransferases in the ATII subfamily that use α-ketoglutarate as the amino group acceptor: the red circle indicates the arginine (R 141 in GABA-T_E.coli) that forms a salt bridge with the γ-carboxylic group of α-ketoglutarate, and the red triangles indicate the glutamate and the arginine of the gateway system. The blue triangle points to the arginine typical of AT-II enzymes that use pyruvate as the amino group acceptor. [4]

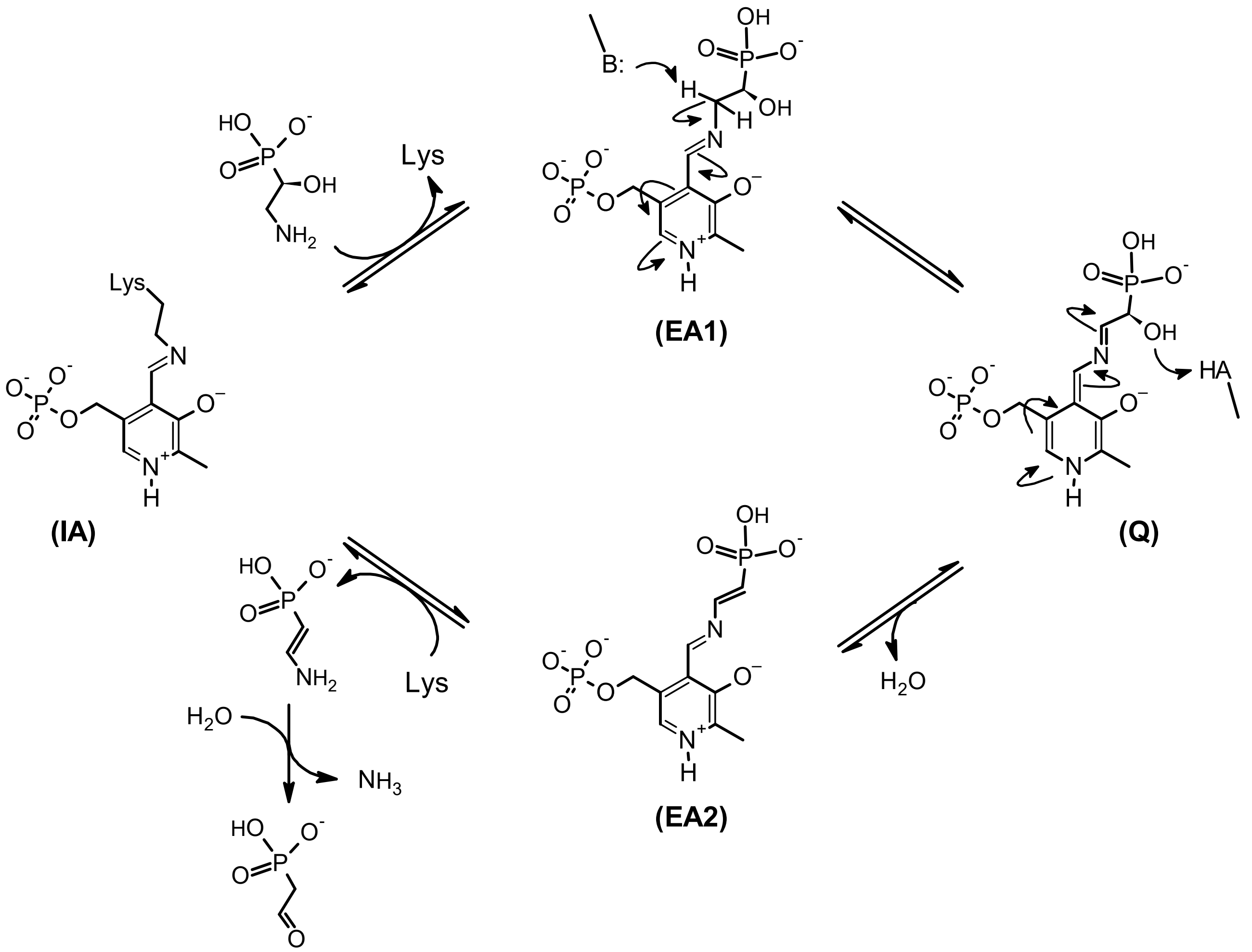

**Figure S2** – Proposed mechanism for the 1,2-elimination reaction catalyzed by PbfA, based on the enzymatic mechanisms of threonine ammonia-lyase and of other PLP-dependent lyases [8–10]. After entering the enzyme active site, *R*-HAEP can attack the Schiff base formed by PLP with the active site lysine (internal aldimine: IA) and – passing through a gem-diamine intermediate (not shown) – complete the transaldimination step to yield the external aldimine (EA1). Proton abstraction from the amino carbon, followed by acid-catalyzed elimination of the OH group, can yield the Schiff base of PLP with ethylene 2-amine phosphonate (EA2). These two latter steps may be concerted, in which case a PLP- stabilized carbanion (quinonoid intermediate; Q) would not form. It is assumed that the eventual hydrolysis of ethylene 2-amine phosphonate (to generate PAA and ammonia) occurs after this compound is released from the active site [8,9].

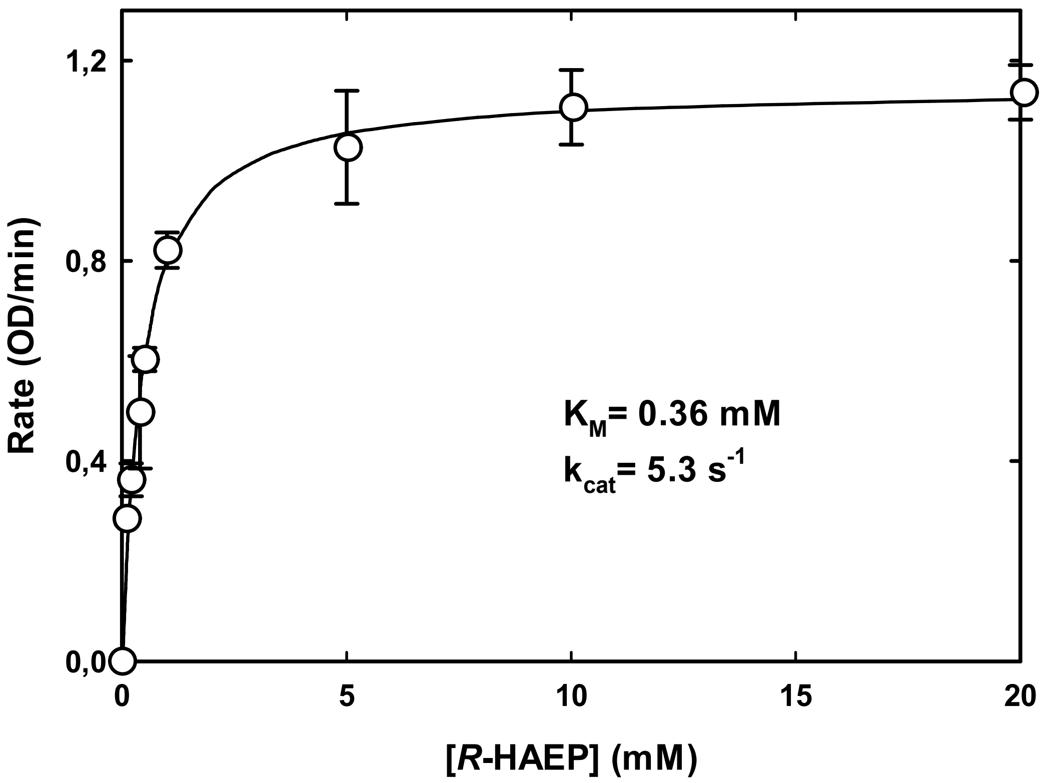

**Figure S3** – Dependence of PbfA activity on the concentration of *R*-HAEP, as measured by the coupled assay with PhnX and ADH in TEA-HCl buffer pH 8.0, 25°C. The reaction mixture contained ~0.25 mM NADH, 5 mM $MgCl_2$, 0,07% BSA, 5 μM PLP, 0.58 μM PbfA, 4 μM PhnX and 9 U ADH.

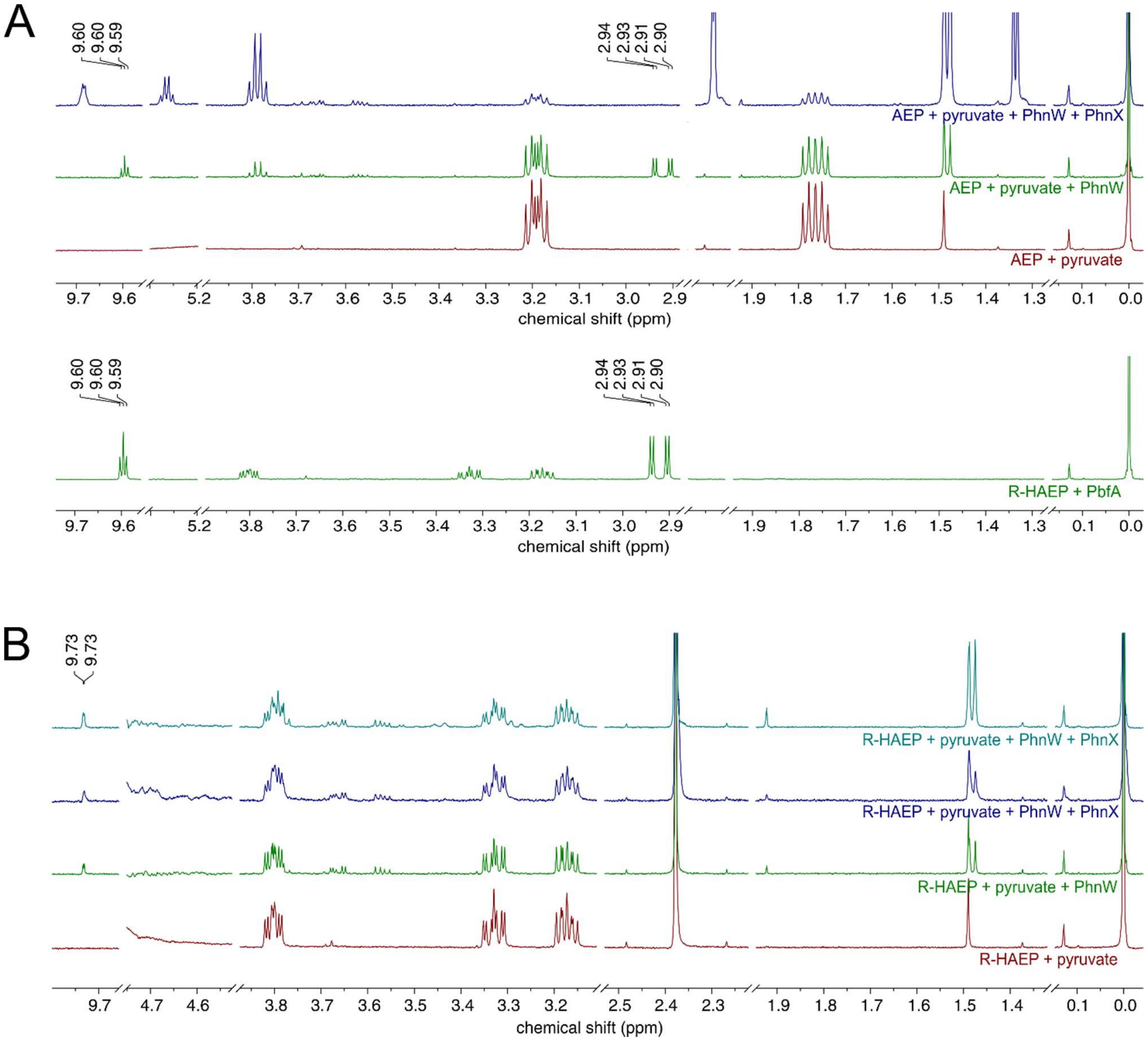

**Figure S4** – $^{1}$H-NMR spectra documenting the reaction of PhnW with AEP and *R*-HAEP. (A) Transamination of AEP. The spectra of a mixture of AEP and pyruvate before (bottom, red line) and after (middle, green line) a 24-min incubation with PhnW are compared. The new peak at 1.47 ppm corresponds to L-Alanine, while the peaks at 2.90-2.94 ppm and at 9.6 ppm correspond to PAA. Addition of PhnX (top, blue line) led to disappearance of PAA and to the appearance of new peaks attributable to acetaldehyde. For comparison, the middle spectrum from figure 4 of the main text (referring to the reaction of PbfA with *R*-HAEP) is shown beneath the spectra referring to the AEP transamination. (B) Reaction of PhnW with *R*-HAEP. The bottom spectrum (red) represents a mixture of *R*-HAEP and pyruvate. After incubating with PhnW for over two hours (green line) some new peaks appeared, corresponding to alanine and to an aldehydic hydrogen from an unknown compound (the expected transamination product, in addition to alanine, would be *R*-1-hydroxyphosphonoacetaldehyde). Addition of PhnX (dark blue line) brought about no substantial changes, but some new peaks started to appear after an extended incubation of the mixture (turquoise line).

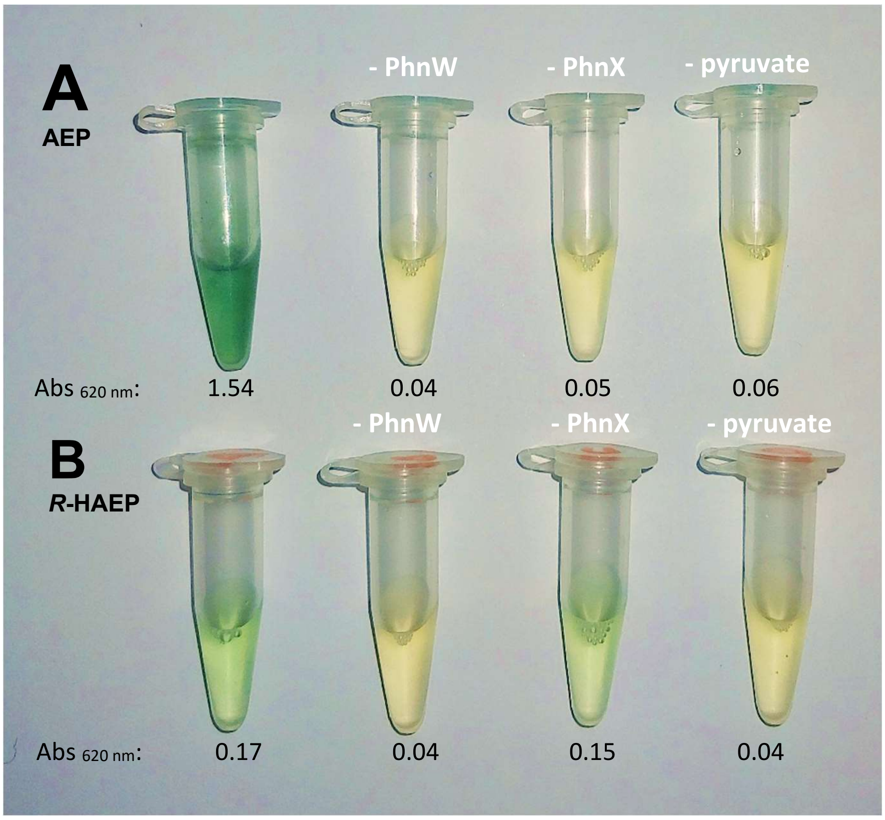

**Figure S5** – Colorimetric detection of phosphate release in the reactions catalyzed by PhnW and PhnY from *V. splendidus* with AEP or *R*-HAEP. (A) Reaction with AEP. The complete reaction mixture (leftmost tube) contained 50 mM TEA pH 8.0, 1 mM DTT, 5 μM PLP, 100 mM KCl, 5 mM $MgCl_2$, 2 μM PhnW, 2 μM PhnX, 1 mM of AEP and 1 mM of pyruvate. The other three tubes show control reactions in which either PhnW, PhnX or pyruvate were omitted. Reactions were carried out at room temperature and they were stopped by adding the BIOMOL® Green reagent after 1 h. (B) Reaction with *R*-HAEP. The experiment was conducted as above, except that *R*-HAEP was used in place of AEP. The greenish color in the first and third tube (but not in the other controls) indicates a weak release of phosphate dependent on the transaminase activity of PhnW, but independent of PhnX.

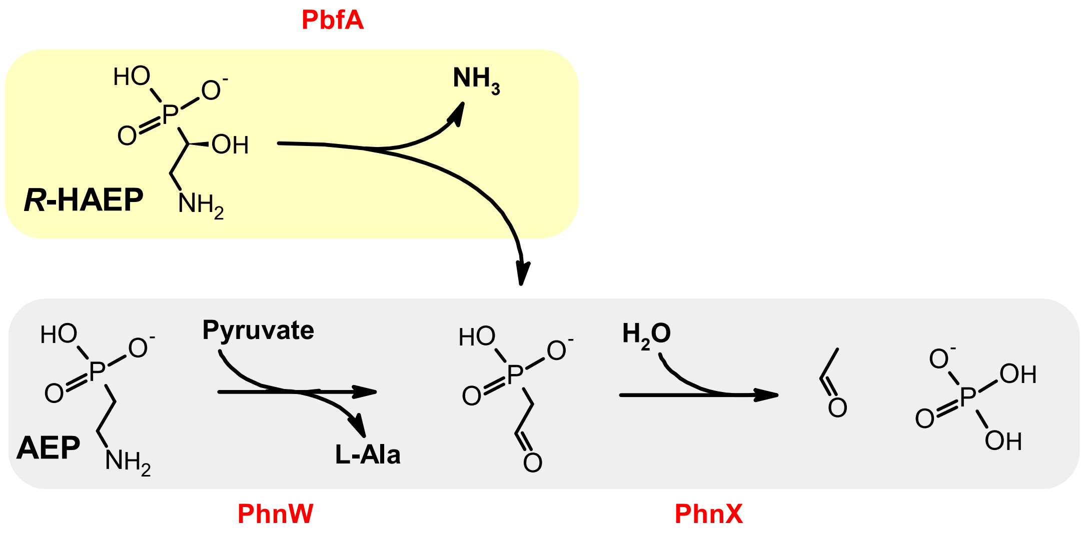

**Figure S6** – A summary of how the PbfA activity (yellow background) integrates into the established PhnW-PhnX catabolic pathway (gray background).
